# Comparative profiling of SARS-CoV-2 variant infections reveals diverse impacts on host cell RNA and RNA binding protein distribution and regulation

**DOI:** 10.64898/2026.01.22.701202

**Authors:** Grégoire De Bisschop, Tram Pham, Nicolas Jacques-De Vries, Monford Paul Abishek Nelson Vimalanathan, Jonathan Boulais, Gabrielle Deschamps-Francoeur, Eunjeong Kwon, Eric Cohen, Eric Lécuyer

## Abstract

RNA viruses perturb cellular RNA metabolism to promote viral transcript expression and genome replication. However, the extent of host cell RNA reprogramming and the consequences on RNA binding protein (RBP) expression and localization remain poorly understood. Here, we utilized an integrated set of transcriptomics, proteomics, and high-content imaging approaches to compare the impact of different SARS-CoV-2 variants, including an early viral isolate and later emerging variants of concern (Alpha, Delta and Omicron) on the host cellular RNA and RBP landscapes. While all variants deeply remodelled the host transcriptome in a subcellular compartment-specific manner, we found that Omicron variants express lower levels of viral RNA products compared to earlier variants, while still driving efficient production of viral proteins. The surge in viral RNA expression correlated with altered subcellular RNA biotype distribution, cytoplasmic mRNA exclusion, the selective disruption of the localization of polyA-tail interacting proteins, and the nuclear accumulation of non-coding transcripts such as RN7SL family members and MALAT-1. Unexpectedly, we found many RBPs that exhibiting altered subcellular localization following abiotic innate immune stimulation with poly(I:C) but remained unaltered in cells infected with SARS-CoV-2 variants, revealing a capacity for these viruses to limit the overall impact on the RBP subcellular distribution. Our results further indicate that Omicron infection sustains higher expression levels of proteins components of the signal recognition particle and translation factors, which may enable a more effective production of immune response and viral components. Altogether, this work highlights the deep impact of SARS-CoV-2 infection on host cell RNA regulatory landscape and provides new insights to explain the differential impact of Omicron compared to early SARS-CoV-2 variants.

**Highlights:**

- SARS-CoV-2 variants drive varying degrees of viral RNA expression in infected Calu-3 cells
- Infection reshapes the host cell RNA landscape in a cell-compartment specific manner
- RN7SL RNA exhibits a strong increase in relative abundance and accumulates in the nucleus of infected cells
- SARS-CoV-2 infections impact the localization of polyA-tail binding proteins while blocking the nuclear accumulation of immune response-sensitive RBPs
- Omicron infections are less disruptive to cellular function and sustain the production of translation factors and components of the SRP complex

## Introduction

RNA viruses are a major source of emerging infectious diseases ^1^. Their inherently high mutation rate favors repeated spillovers and immune surveilance escape, despite acquired immunity and prophylactic measures. For instance, several SARS-CoV-2 variants of concern (VOCs) with devastating effects emerged within a few years and still pose a threat to human health. SARS-CoV-2 variants carry mutations allowing them to escape detection by acquired immune pathways, permitting them to thrive via alternative molecular mechanisms.

SARS-CoV-2 is a positive-strand, single-stranded RNA virus, for which the ∼30 kb genomic RNA possesses typical features of a host mRNA, such as the 5’ cap and a 3’ poly(A) tail ^2^. Thus, once delivered to the cytoplasm of target cells, it is translated by the host machinery to yield two polyproteins, ORF1a and ORF1ab. Proteolytic processing of these polyproteins generates a set of non-structural proteins that assemble to form a replication and transcription complex. This complex generates negative-sense copies of the genomic RNA as well as numerous shorter, subgenomic RNA through discontinuous transcription. Viral RNA synthesis occurs in viral factories, localized within ER-derived double-membrane vesicles (DMVs), and these transcripts are further processed through capping and polyadenylation. Although SARS-CoV-2 encodes its own capping activity ^3^, it is not known how polyadenylation occurs. Translation of the subgenomic mRNA produces structural proteins that assemble new viral particles. In addition to these functions that directly promote replication, viral proteins interfere with many host proteins and RNAs to take control over host machineries ^4^. Viral RNAs have also been shown to interact with numerous host proteins ^5^ and RNA molecules ^6,7^. These SARS-CoV-2 RNA interacting proteins are known to localize to various subcellular compartments, although SARS-CoV-2 replication occurs solely in the cytoplasm.

RNA binding proteins (RBPs) represent a particularly important class of cellular regulatory factors that modulate RNA virus infection cycles. These factors, of which >1,500 are encoded in the human genome, play essential roles in controlling various aspects of RNA mutation, function and fate within intracellular space. Several RNA viruses have been shown to disrupt the subcellular localization properties of RBPs. For instance, Sindbis virus induces the relocalization of DDX1 and GEMIN5 to viral replication factories ^8^. The surge in viral RNA abundance, combined with a decrease in host RNA levels, is thought to rewire RBPs toward viral RNA binding ( DOI: 10.1016/j.molcel.2019.01.017). In addition, pervasive cytosolic RNA degradation, either mediated by RNAse L or viral endonucleases, such as SARS-CoV-2 NSP1, has been shown to trigger a broad nuclear accumulation of RBPs, most prominently the poly(A)-binding protein PABPC1 ^9,10^.

To gain deeper insights into SARS-Cov-2 infection properties, we sought to comparatively evaluate the extent to which infections with different SARS-CoV-2 variants impact host transcriptome regulation, as well as the subcellular distribution properties of RBPs. To this end, we compared the transcriptomic profiles of Calu-3 cells infected with pre-VOC, Alpha, Delta and Omicron SARS-CoV2 variants, revealing a striking reduction in the expression of cytoplasmic poly-A transcripts and an over-abundance of nuclear-enriched RNA biotypes across all infections contexts. Intriguingly, while Omicron infected cells exhibited similar levels of viral nucleocapsid and spike proteins, as assessed by imagining, they expressed lower levels of viral genomic and sub-genomic RNAs compared to the other variants. We also conducted comparative systematic immunofluorescence (IF)-based screens, using a collection of ∼185 validated RBP antibodies, in cells that were either treated with poly(I:C) to stimulate the innate response or infected with SARS-CoV-2 variants. Strikingly, of the 19 RBPs that exhibited altered localization in response to poly(I:C) treatment, the majority were not impacted in the infection context, except for poly-A binding factors. Finally, in addition to expressing higher levels of immune response proteins, we found that cells infected with Omicron exhibited higher levels of factors involve in mRNA translation, such as components of the signal recognition particle (SRP), translation initiative factors and ribosomal proteins.

## Results

### Viral RNA load differs across SARS-CoV-2 variants

SARS-CoV-2 infection, or the expression of individual viral proteins in isolation, has been shown to perturb cellular post-transcriptional gene regulatory pathways ^4,5,11^. To investigate the impact of different SARS-CoV-2 variants on RNA-regulatory machineries, we infected Calu-3 cells with a Wuhan-Hu-1-like isolate (pre-VOC) or three variants of concern (VOCs): Alpha, Delta and Omicron BA.1. Calu-3 cells are naturally permissive to early SARS-CoV-2 variants and Omicron subvariants ^12^, making them a suitable model system. These SARS-CoV-2 variants carry several missense mutations, particularly within the region encoding the Spike protein (**Figure S1A**). To achieve a high percentage of infected cells with minimal cytotoxicity, we optimized the multiplicity of infection (MOI) and time parameters. The selected MOI’s ranged from 0.01 for Omicron to 0.1 for the pre-VOC strain, conditions that yield comparable infection rates at 24 and 48 hours post-infection (hpi), ranging from 45-60% of infected cells, as determined by FACS-labeling using an anti-Nucleocapsid antibody ^13^ (**Figure S1B**). Replicate cellular samples were fixed for imaging or processed for RNA and protein extraction at 48 hpi to maximize the percentage of infected cells (**Figure 1A**), as done previously ^14^.

**Figure 1.**
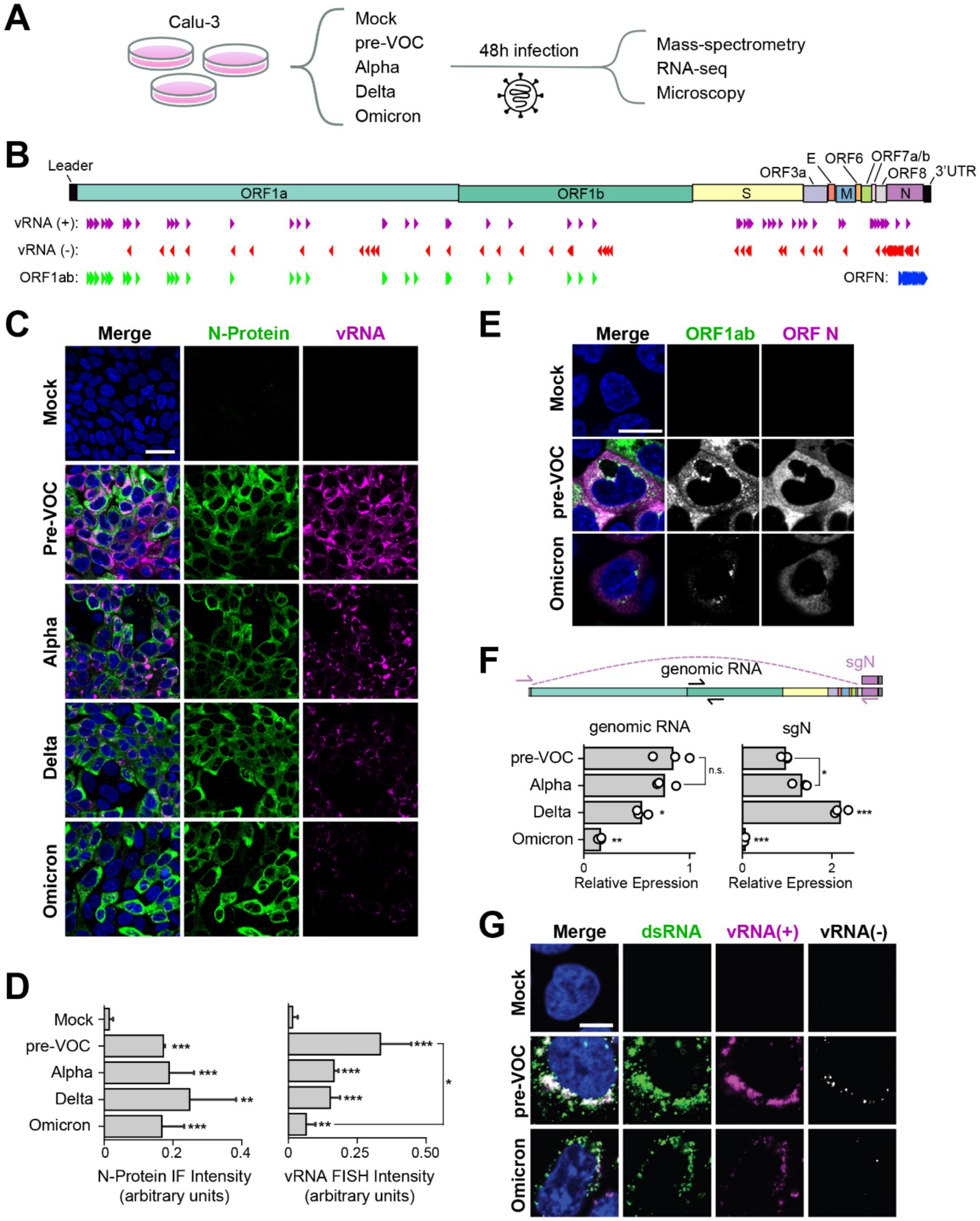
RNA viral load varies across SARS-CoV-2 variants. **A)** Schematic of the experimental workflow used in this study. **B)** Architecture of the SARS-CoV-2 genome, with binding sites of FISH probe sets indicated by color triangles. **C)** Immunofluorescence staining of infected Calu-3 cells with an antibody against SARS-CoV-2 nucleocapsid protein (N-protein) combined with viral RNA smiFISH. Scale bar: 40 µm. **D)** Mean of total viral RNA FISH and nucleocapsid IF intensities per infected cell, as shown performed in (**C**), with standard deviation across n=3 replicates**. E)** Viral RNA smiFISH conducted on control or infected Calu-3 cells using the indicated probes specific targeting the ORF1ab (green) or ORF-N (magenta) sub-genomic regions. Scale bar: 20 µm. **F)** Location of RT-qPCR primers used to assess ORF1ab and ORF N sub-genomic RNA expressions and corresponding RT-qPCR 2^-ΔΔCt^ results using pre-VOC as the reference. Data represents n=3 replicates. Statistical significance is denoted as follow: * p<0.05, ** p<0.005, *** p<0.0005. **G)** Viral RNA smiFISH targeting positive-sense (+) and negative-sense (-) viral RNA, combined with immunofluorescence co-labeling with the 9D5 antibody detecting dsRNA structures. Scale bar: 10 µm.

We first assessed viral RNA load in infected cells using single molecule inexpensive RNA *in situ* hybridization (smiFISH) with different probe sets (**Figure 1B**), including: ***i)* vRNA(+)** antisense probes covering the entire length of the SARC-CoV2 genomic sequence, expected to detect genomic RNA (gRNA) and most subgenomic (+) strand RNA (sgRNA), ***ii)*** a distinct set of probes in sense orientation, designated **vRNA(-)**, allowing detection of (-) strand copies of the viral genome; and ***iii)*** antisense probes covering the **ORF1ab** or **ORF-N** regions capable of detecting (+) strand genomic RNA and subgenomic RNAs derived from the corresponding regions ^15^. Of note, probes had on average more than 99% sequence identify across variants (note shown). Strikingly, while immunofluorescence (IF) against the nucleocapsid (N) protein or spike protein showed comparable staining intensities (**Figure 1C** and **S1C**), smiFISH with the vRNA(+) probe set revealed a progressive decreased in detection levels across variants, with Omicron BA.1 exhibiting significantly lower labeling compared to the earlier variants (**Figure 1C-D**). This low expression was also apparent in infections with several Omicron subvariants, including BA.1-2, BA.5, and JN.1 (**Figure S1D**). Consistent with the vRNA (+) labeling, double-smiFISH with ORF1ab and ORF-N probes also revealed a reduced expression signal in Omicron infected cells versus pre-VOC (**Figure 1E**). Notably, these probes gave distinctive labeling features, with ORF1ab exhibiting a perinuclear foci distribution pattern, likely corresponding ER-associated viral replication organelles, while ORF-N probes produced a more diffuse cytoplasmic signal. To validate these observations, RT-qPCR was performed using oligonucleotides designed to amplify gRNA (targeting a region with ORF1b) or the ORF-N sgRNA (**Figure 1F**), revealing that gRNA expression progressively decreases across variants, whereas ORF-N sgRNA levels increase through Delta, but are strongly decreased in Omicron, consistent with our smiFISH data. Of note, RT-qPCR values were normalized to 18S rRNA as the endogenous control, as its abundance remains constant across samples according to bioanalyzer traces (**Figure S1E**) and as previously observed ^11,16^. Finally, double smiFISH with vRNA(+) and vRNA(-) probe sets, in conjunction with IF using the 9D5 antibody that recognizes double stranded RNA (dsRNA) structures, revealed that negative-stranded viral RNA is only weakly detected within disparate cytoplasmic foci, both in pre-VOC and Omicron infected cells, generally occurring within much broader territories of vRNA(+) signal (**Figure 1G**). Strikingly, the dsRNA antibody gave an intense cytoplasmic labeling pattern in infected cells, which partially colocalized with positive strand RNA, while the antibody signal was negligeable in uninfected cells. This suggests that SARS-CoV-2 infection drives the accumulation of dsRNA structures in the cytoplasm of infected cells, structures that occur both within vRNA-rich territories labeled with vRNA(+) probes and within other regions of the cytoplasm.

### Host RNA landscape following SARS-CoV-2 infection

To assess the impact of SARS-CoV-2 variant infections on host cell gene expression, we next sequenced the transcriptomes of mock and infected Calu-3 cells via Illumina-seq with ribosomal RNA depletion (RD). As expected, a high proportion of reads mapped to the viral genome, consistent with prior reports ^17^, ranging from 80-90% of total reads in pre-VOC, Alpha and Delta specimens, and 50% of reads for Omicron samples (**Figure 2A**). Principal component analysis (PCA) revealed the distinctive segmentation of mock and omicron samples, with pre-VOC, Alpha and Delta specimens co-clustering together (**Figure S2A**). Our expression data for pre-VOC were well correlated with previous data generated with Calu-3 cells and a similar SARS-CoV2 strain (**Figure S2B**) ^18^. Consistent with the predominance of viral reads in sequencing libraries, we noted a 2-to 10-fold reduction in the number of detected host transcript in infected samples, depending on the expression threshold used (**Figure 2B**). A similar trend was observed for non-coding RNA classes, with a subset of these transcripts being over-represented in infected cells relative to mock samples (**Figure S2C**). RNA biotype analysis revealed a decrease in mRNA reads in infected samples, consistent with SARS-CoV-2 induced mRNA degradation, while several classes of non-coding RNAs were increased, in particular lncRNAs, miscRNA and snRNA (**Figure 2C**). Interestingly, when viral RNA is considered among coding transcripts, the total coding content in sequencing reads was similar or increased in infection specimens (**Figure 2D**).

**Figure 2.**
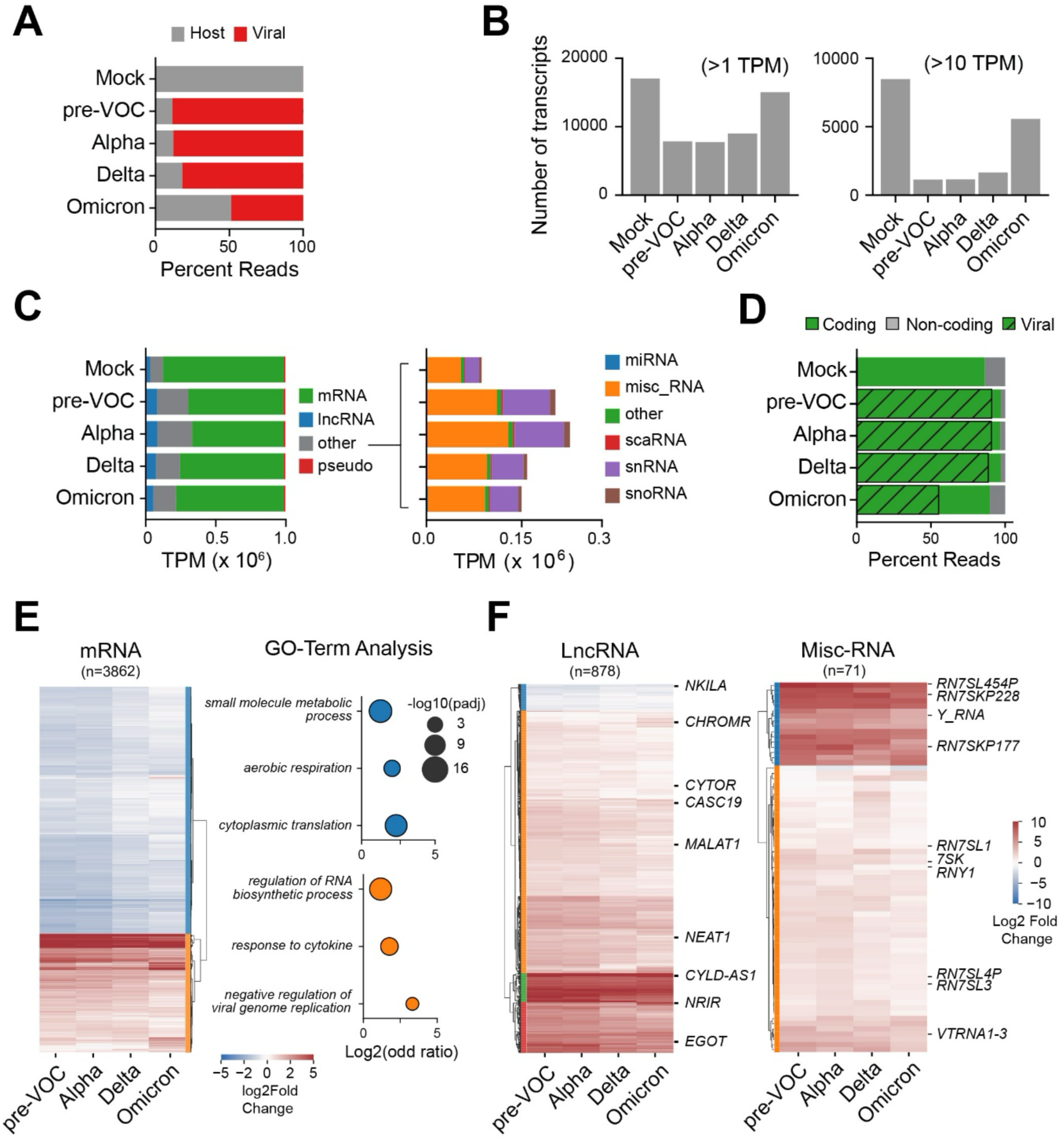
Impact of SARS-CoV-2 infections on the host cell transcriptome. **A)** Average proportion of viral and host sequencing reads in uninfeced Calu-3 cells (mock) and in cells infected with pre-VOC, Alpha, Dela or Omicron viral strains for 48h, assessed via Illumina ribo-depletion RNA-sequencing. **B)** Number of transcripts detected across samples using TPM cutoffs of 1 (top) or 10 (bottom). Proportions of unchanged, up- and down-regulated transcripts are indicated in gray, red and blue respectively. **C)** <u>Left</u>: Transcripts per million (TPM) distribution of the major expressed RNA biotypes (i.e. mRNA, lncRNA and pseudogenes) in mock versus SARS-CoV-2 infected Calu-3 cells. <u>Right</u>: Detailed TPM distribution for minor RNA biotypes composing the ‘other’ category in the left side panel. **D)** Percentage of reads mapping to coding or non-coding genes. Viral reads are indicated by hatched lines. **E)** <u>Left</u>: Differential expression analysis of protein coding mRNAs across all variants relative to mock specimens, revealing a cluster of mRNAs that are under-represented (blue, n=2611) or over-represented (orange, n=1251) in infected cells relative to mock specimens. <u>Right</u>: Bubble plots conveying gene ontology (GO) terms that are statistically enriched within the blue and orange mRNA clusters shown in the left panel. **F)** Differential expression analysis of lncRNA (left) and miscellaneous RNA (right).

Comparing the total number of transcripts detected in mock versus all infection samples combined revealed a common set of 12,416 detected transcripts (TPM≥1), with 2,699 and 614 being exclusively detected in mock or infected specimens, respectively (**Figure S2D-E**). Differential expression analysis identified 3,862 mRNAs that are significantly over- or under-represented across variants (**Figure 2E**, left panel), with over-represented mRNAs being enriched for immune response-related Gene Ontology terms (**Figure 2E**, right panel), while the under-represented class having links to translation and respiration. By contrast, most differentially expressed lncRNA and miscRNA were over-represented in infection samples (**Figure 2F**), including several lncRNA known to be induced by viral infections and cellular stress responses (e.g. *EGOT, NRIR, CHROMR*) ^19–23^.

### SARS-CoV-2 infection differentially impact subcellular transcriptomes

To characterize transcriptomic changes that may be occurring at the subcellular level in SARS-CoV-2 infected cells, we next contrasted our transcriptomic signatures to previously published cell fractionation and RNA-seq (CeFra-seq) datasets ^24^. This previous work characterized RNA populations enriched in nuclear, cytosolic, cytoplasmic-insoluble (cytoskeletal) and endomembrane fractions of HepG2 cells. Strikingly, this analysis revealed that reads corresponding to mRNAs known to be enriched in cytoplasmic compartments (e.g. cytosolic, cytoskeletal and endomembrane) were depleted in infected samples relative to mock, while reads of nuclear-enriched mRNAs and lncRNAs were increased (**Figure 3A**). Additionally, reads corresponding to cytosolic miscRNAs accumulated in infected samples. When considering individual transcripts, contrasting the steady-state subcellular distribution features in CeFra-seq data versus their expression parameters in infected Calu-3 cells (**Figure S3A**), we found that normally abundant cytoplasmic mRNAs (e.g. *RPS12, RPL35, GAPDH, YBX1*) are depleted upon infection. By contrast, several stress response-associated mRNAs that are enriched in cytoskeletal (e.g. *OAS2, IRF2, IFIH1, STAT1*) and nuclear (e.g. *SPEN, NFKBIZ, JUN*) compartments generally showed an enhanced relative detection in infected cells. We also noted a general increase in expression of non-coding lncRNA (e.g. *MALAT-1, NEAT1, CYTOR*), miscRNA (e.g. *RN7SL, RNY1, SNORD3A, SCARNA7-10*) and snRNAs (*NRU1*), regardless of their steady-state subcellular distribution (**Figure S3A**). These findings are consistent with the documented impact of infection on cytoplasmic mRNA degradation driven by RNAse L activation and Nsp1 activity ^9,25^. By contrast, the increased relative detection of reads for typically nuclear-enriched RNA biotypes in infected cells likely reflects their shielding from virus-induced cytoplasmic degradation mechanisms, blockage of RNA nuclear export, or both ^25,26^.

**Figure 3.**
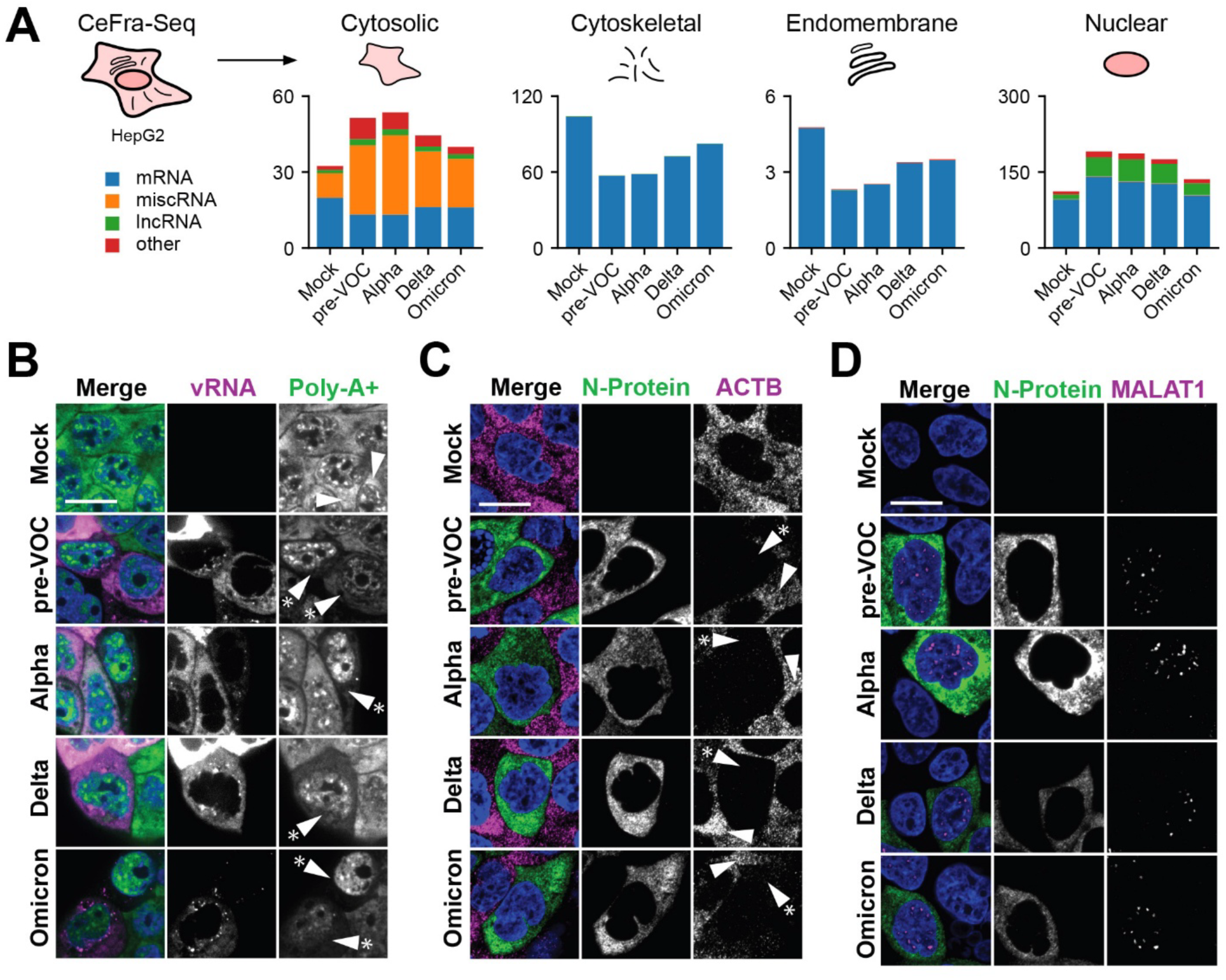
SARS-CoV-2 infections selective impact host subcellular transcriptomes. **A)** Top: Schematic of subcellular compartments analyzed in CeFra-seq datasets previously generated with HepG2 cells. RNA purified from cytoplasmic (i.e. cytosolic, endomembrane and cytoplasmic insoluble/cytoskeletal) and nuclear compartments was subjected to RNA-sequencing, identifying collections of transcripts enriched within specific subcellular compartments ^24^. Bottom: Comparative analysis of summed TPM values of previously defined compartment-enriched transcripts within mock or infected Calu-3 cells. Host transcripts are binned by subcellular localization. **B)** FISH-based co-detection of polyA (+) RNA combined with viral RNA FISH in mock or SARS-CoV-2 infected Calu-3 cells. **C-D)** RNA FISH of ACTB mRNA (**C**) and MALAT1 lncRNA (**D**) combined with N-protein protein immunofluorescence. For **B-C**, the arrow heads indicate uninfected cells, while arrow heads with asterisk point to infected cells. Note the selective reduction in signal for polyA (+) in the cytoplasm of infected cells (**B**), the striking loss of ACTB mRNA signal in infected cells (**C**), and the persistence of MALAT1 nuclear labeling in infected cells (**D**).

We next sought to complement our sequencing-based observations with direct visualization of specific RNA species by smiFISH. While detection of poly-A+ RNA with an oligo-T probe in mock cells displayed a combination of cytoplasmic diffuse and punctate nuclear structures, the cytoplasmic labeling was strongly reduced in infected cells that were positively labeled for vRNA across all SARS-CoV-2 variants interrogated (**Figure 3B, S3B**). The overall poly-A+ RNA intensity decreased in infected cells and, to a lesser extent, in bystander cells (**Figure S3C**). Since the oligo-T probe is expected to detect both host poly-A+ RNA and viral RNA, which has short poly-A tails ^17^, some persistence of poly-A signal in the cytoplasm of infected cells was expected. Strikingly, smiFISH detection of cytoplasmic mRNAs, such as *ACTB* or *EIF4A1*, revealed a near complete loss of signal in infected cells (**Figure 3F, S3D**). By contrast, the signals for nuclear enriched non-coding RNA, such as RNU2 snRNA (**Figure S3E**) and the MALAT-1 lncRNA (**Figure 3D**), or of mitochondrial RNA (**Figure S3F**), were unchanged or appeared more intense in infected cells.

### Infection triggers RN7SL1 nuclear accumulation

We next sought to ascertain which transcripts accounted for the altered expression of cytosolic-associated mRNA and miscRNA in infected cells (**Figure 3A**). Further analysis showed that the top 10 transcripts accounting for the reduction in mRNA reads were coding transcripts that are normally highly expressed, including *S100A6*, *GAPDH*, and ribosomal protein mRNAs (**Figure 4A**, lower panel). By contrast, the increased abundance of cytosolic miscRNA reads was attributable to the specific subsets of small non-coding RNAs, most prominently transcripts of the RN7SL family, the RNA component of the signal recognition particle (SRP) ribonucleoprotein complex implicated in the recruitment and localized translation of mRNAs encoding secretory proteins at the ER, followed by *RNU1* and *RN7SK* family members (**Figure 4A**, upper panel). To further validate these findings, we next employed RT-qPCR analysis (**Figure 4B**). Since Alu repeat elements, the most highly abundant transposons in the human genome, derive from *RNSL7* RNA, we designed probe sets that only amplify *RN7SL* (**Figure 4B**, upper left cartoon). Using two distinct *RN7SL*-selective probe sets, we found that RN7SL1 expression was unaltered during infections across all variants (**Figure 4B**, lower graphs). By contrast, using exon-exon versus exon-intron junction spanning primer pairs to distinguish unprocessed versus mature mRNA, we found that the expression of mature *GAPDH* or *SOD1* mRNAs was strongly reduced in infected cells, while the expression of pre-mRNAs transcribed from both genes was unaffected or even increased in infected cells (**Figure 4C**).

**Figure 4.**
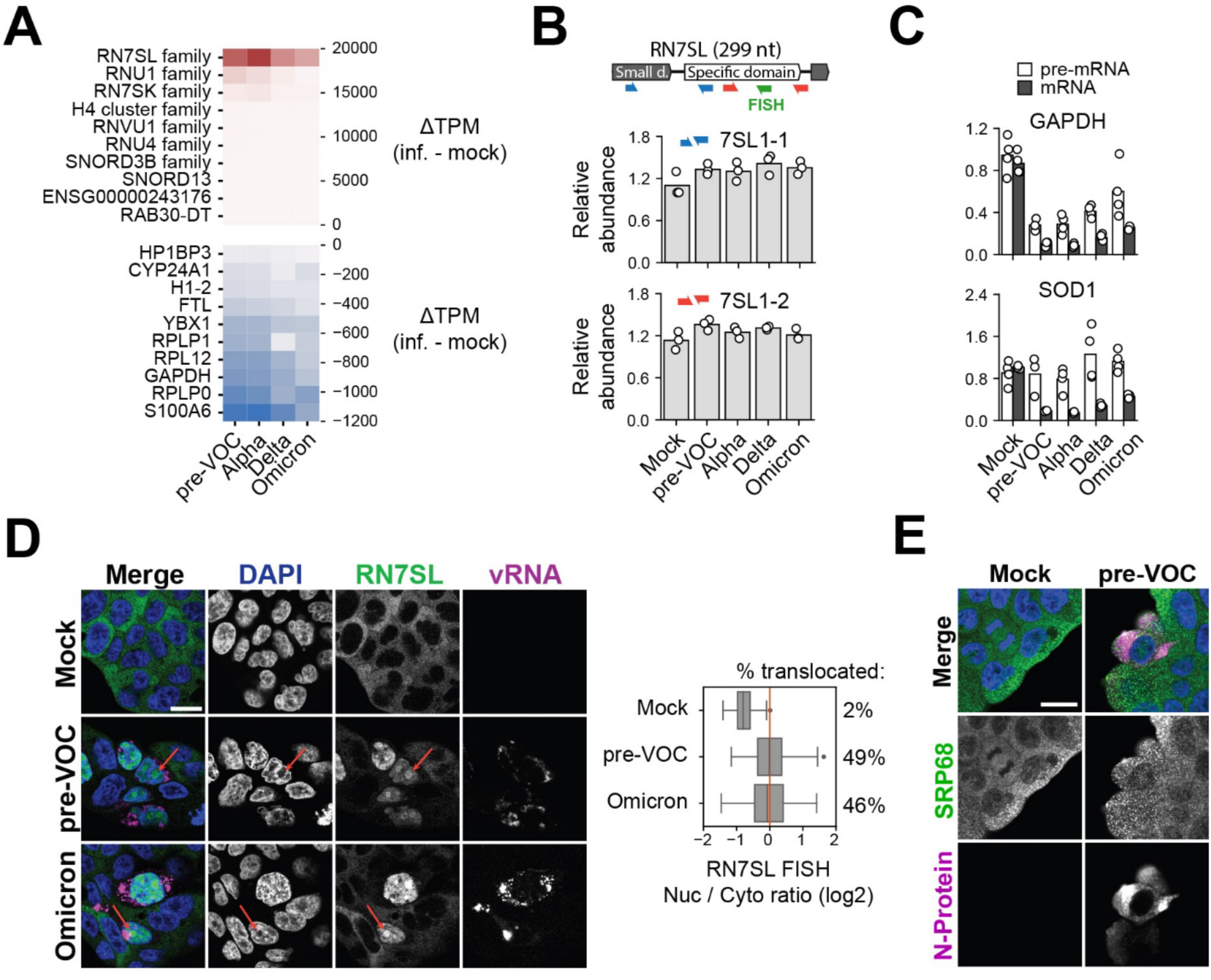
RN7SL1 accounts for the increased miscRNA expression in infected cells and becomes nuclear enriched upon infection. **A)** Top 10 transcripts annotated as cytosolic ranked by the greatest (top) or lowest (bottom) TPM difference between mock and infected conditions. **B)** RT-qPCR analysis for the *RN7SL1* of mock or infected Calu-3 RMA specimens, evaluated using the RN7SL1-specific primer pairs (red and blue) indicated the cartoon. Small d.: small domain. **C)** RT-qPCR analysis of *GAPDH* and *SOD1* mRNA and pre-mRNA relative abundance in mock and infected cellular specimens. **D)** <u>Left:</u> RNA FISH of RN7SL1 for mock, pre-VOC and Omicron infected Calu-3 cells, revealing a strong nuclear enrichment of RN7SL in infected cells. Right: Boxplot showing the log2 ratio of nuclear over cytoplasmic mean intensity in mock vs infected cells. **E)** Immunofluorescence labeling of mock or pre-VOC infected Calu-3 cells with antibodies against SRP68 and N-protein.

Strikingly, when conducting smiFISH using a probe targeting the specific domain of RN7SL in mock specimens versus cells infected with pre-VOC or Omicron, we observed that the normally predominant cytoplasmic localization pattern of RN7SL1 was altered to a striking nuclear distribution in infected cells (**Figure 4D**). This observation is reminiscent of a recent study uncovering a nuclear function for RN7SL1 in response to thermal stress, where it bins to genes repressed by heat shock ^27^. However, by contrast to this previous study, we found that the cytoplasmic distribution pattern of protein components of the SRP, such as SRP68 ^28^, was unaltered in infected cells (**Figure 4E**). This suggests that RN7SL nuclear accumulation may result from a blockage to nuclear export imposed by the infection context, rather than a nuclear recruitment of intact SRP complexes. Overall, the nuclear accumulation of 7SL RNA likely defines an additional mechanism that contributes to the reduced translation of mRNAs encoding membrane-anchored and secretory proteins in SARS-CoV-2 infected cells.

### Selective RBP subcellular localization changes upon infection

Given the extensive transcriptomic alterations induced by infection, we next wanted to study the effects of SARS-CoV-2 variants on the subcellular localization properties of RBP machineries. To distinguish the impact of acute innate immune response activation versus *bona fide* viral infections, we compared the effects abiotic immune stimulation via polyinosinic:polycytidylic acid (polyI:C) transfection, a synthetic dsRNA polymer that activates pattern recognition receptor signaling ^29^, to infections with SARS-CoV-2 variants (**Figure 5A**). Poly(I:C) induces cytoplasmic RNA degradation via RNase L activation and blocks mRNA export, leading to nuclear re-localization of some RBPs and formation of cytoplasmic RNase L-dependent bodies (RLBs) ^30^. Consistent with prior studies ^9^, we found that poly(I:C) transfection caused nuclear accumulation of poly-A+ RNA and the formation of cytoplasmic poly-A+ RLBs (**Figure 5B)**. Poly(I:C) also induced the degradation of cytoplasmic RNAs (e.g., ACTB), while the smiFISH signal for nuclear-enriched transcripts such as Malat-1 is comparable to control cells (**Figure S4A**). The P-body component DDX6 formed fewer, larger granules upon SARS-CoV-2 infection (**Figure S4B**) and poly(I:C) transfection (**Figure S4C**), consistent with stress-response-induced disruption ^31^.

**Figure 5.**
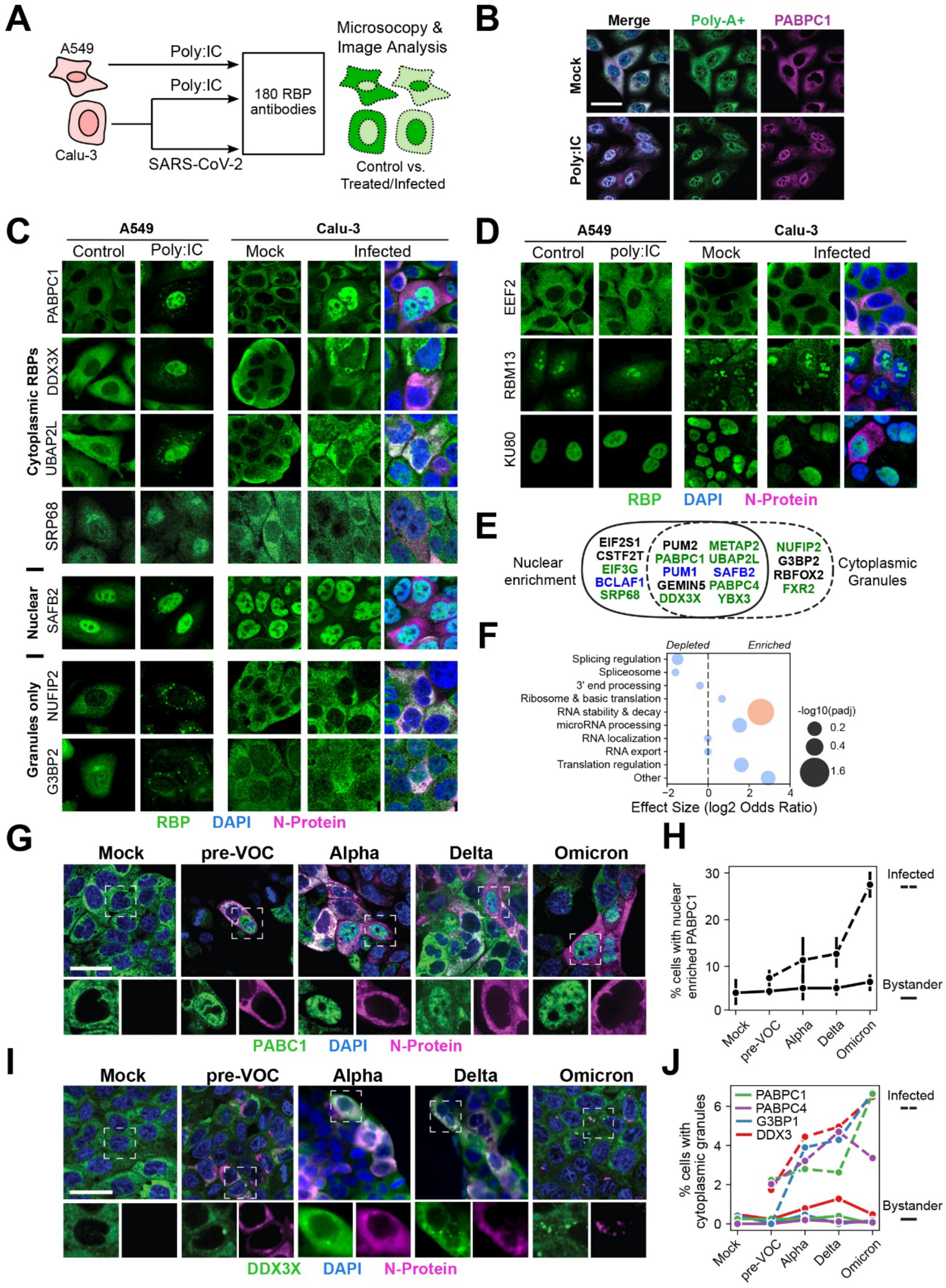
Impact of SARS-CoV-2 infections on the subcellular distribution features of RNA binding proteins. **A)** Schematic of the experimental workflow. A549 and Calu-3 cells were transfected with poly:(IC) for 6 hours. Calu-3 cells were used for infection with SARS-CoV-2 pre-VOC for 48 hours. Cells were then fixed, immuno-labelled with 188 different RBP antibodies and analyzed by high-content microscopy to identify subcellular localization changes. **B)** poly(A) RNA FISH combined with PABPC1 immunofluorescence in control and poly:(IC) transfected cells. **C)** Examples of RBPs with altered subcellular localization either upon poly:(IC) transfection in A549 (left) or SARS-CoV-2 infection in Calu-3 (right). **D)** RBPs with different subcellular localization patterns that are unaffected by poly:(IC) transfection. **E)** Venn diagram showing the intersection of RBP forming cytoplasmic granules (left) or relocalizing to the nucleus (right) upon poly:(IC) transfection. Green: RBPs primarily cytoplasmic, blue: RBPs primarily nucleoplasmic. **F)** Functional annotations enrichment analysis among RBP with perturbed subcellular localization. **G-J)** Immunofluorescence staining of PABPC1 (**G**) or DDX3X (**I**) upon infection with different variants. Percentage of uninfected (solid line) and infected (dashed line) cells with nuclear enriched PABPC1 (**H**) or cytoplasmic granules (**J**).

We next conducted systematic high-content immunofluorescence (IF) screens using a collection of 185 validated antibodies targeting human RBPs ^32,33^. Specifically, we performed comparative IF assays of A549 and Calu-3 cells transfected with poly:IC, or untransfected cells as controls, as well as Calu-3 cells infected with the pre-VOC or mock as control. A549 cells were included for comparison since they are more efficiently transfected with poly(I:C) and much easier to image compared to Calu-3 cells. Approximately ∼10% of RBPs (19 of 185 antibodies tested) showed altered localization after poly(I:C) transfection, primarily exhibiting a shift to the nucleus or cytoplasmic granules (**Figure 5C**, left; **Figure S4D**) while the majority of RBPs remained unaffected by either poly(I:C) or pre-VOC infection (**Figure 5D**). These RBPs were predominantly cytoplasmic under normal conditions (**Figure 5E**). Beyond PABPC1 and PABPC4 nuclear translocation, we identified novel poly(I:C)-responsive RBPs, including the known stress granule proteins DDX3X and UBAP2L, as well as SRP68. Functional analysis of poly(I:C)-responsive RBPs revealed enrichment for “RNA decay and stability” (FDR < 0.05) (**Figure 5F**) using a curated annotation list ^34^. Poly:IC induced similar effects in Calu-3 cells (**Figure S4E**).

Surprisingly, SARS-CoV-2 variant infections only induced the altered distribution of a small subset of proteins, most prominently the nuclear accumulation PABPC1 and PABPC4 (**Figure 5C** and **S4D**). Also, while RBPs such as DDX3X and UBAP2L localized to cytoplasmic granules in a subset of bystander cells, they were not redirected to the nucleus during infection, unlike poly(I:C) transfection conditions. Overall, the distribution patterns of most poly(I:C)-sensitive RBPs remained unaltered upon SARS-CoV-2 infection. To assess variant-specific effects, we examined PABPC1 nuclear translocation across variants and found that Omicron infection induced PABPC1 nuclear translocation twice as frequently as other variants, correlating with its lower viral RNA load (**Figure 5G-H**). DDX3X, which translocates to the nucleus with poly(I:C) but not pre-VOC infection, also showed no nuclear translocation with VOCs (**Figure 5I**). Infection induced cytoplasmic granule targeting of RBPs, such as DDX3X, G3BP2, PABPC1 and PABPC4 in a small percentage of cells, with Omicron showing a higher frequency of this phenotype versus earlier variants (**Figure 5J**), as previously reported ^35^.

The absence of nuclear translocation of poly(I:C)-sensitive RBPs in the cytoplasm of infected cells may result from disrupted nucleo-cytoplasmic trafficking ^4^ or direct interaction with cytoplasmic by viral components. To address this hypothesis, we cross-referenced the list of poly(I:C)-sensitive RBPs with viral RNA interactors and viral proteins interactors (data not shown). We found that, out of 18 poly(I:C)-sensitive RBPs present in all datasets, 6 (33%) were known direct interactors of SARS-CoV-2 RNA as determined by RAP-MS ^36^ (**Figure S4F**) and 4 (22%, p-value) maintained protein-protein interactions (PPI) with viral proteins ^37^ (**Figure S4G**). This is significantly more than expected by chance (p-value = 0.001 and 0.003, respectively, Fisher exact test). Moreover, several of these RBPs (e.g. DDX3, G3BP2, PABPC1) were shown to contribute to viral replication ^38,39^. These results suggest that SARS-CoV-2 prevents nuclear translocation of RBPs following viral sensing, potentially through direct interactions.

### Evaluating the impact of SARS-CoV-2 infection on protein expression and translation

To gain additional insights into the impact of SARS-CoV-2 variant infections, we also conducted parallel proteomic profiling via liquid chromatography tandem mass spectrometry (LC-MS/MS) on infected Calu-3 specimens versus mock as a control, using four replicates per condition. Infected samples gave distinctive signatures compared to mock based on principal component analysis (**Figure S5A**). More than 4,000 proteins were detected across each sample type, among which only a small subset were specific to mock (118) or infected (621) specimens (**Figure 6A** and **S5B**). As described previously by Krogan and colleagues ^14^, we found that Omicron BA.1 infection induced the expression of innate immune response factors, as opposed to pre-Omicron variants (**Figure 6A**, **S5C**, **S6A**). Comparative analysis of RNA and protein differential expression changes revealed a lack of correlation for pre-VOC, Alpha and Delta specimens, while they were moderately correlated (R=0.46) for Omicron (**Figure 6B**). This is expected from the known inhibition of translation caused by infection, and perhaps more generally because of RNA expression buffering by translation. However, this effect was less marked in Omicron. While most differentially expressed coding genes were downregulated at the RNA level and unchanged at the protein level, we observed a higher proportion of genes being upregulated at both levels in Omicron specimens (**Figure 6C**). This suggests that translation is sustained more efficiently following Omicron infection, at least for a subset of genes. Consistent with this notion, closer evaluation of protein intensities across cellular specimens revealed a specific increase in expression of components of the SRP complex and proteins associated with mRNA translation (e.g. ribosomal proteins, translation initiation factors) in Omicron infected cells (**Figure 6D**, **S6B**), while the expression of splicing associated proteins was more variable (**Figure S6C**).

**Figure 6.**
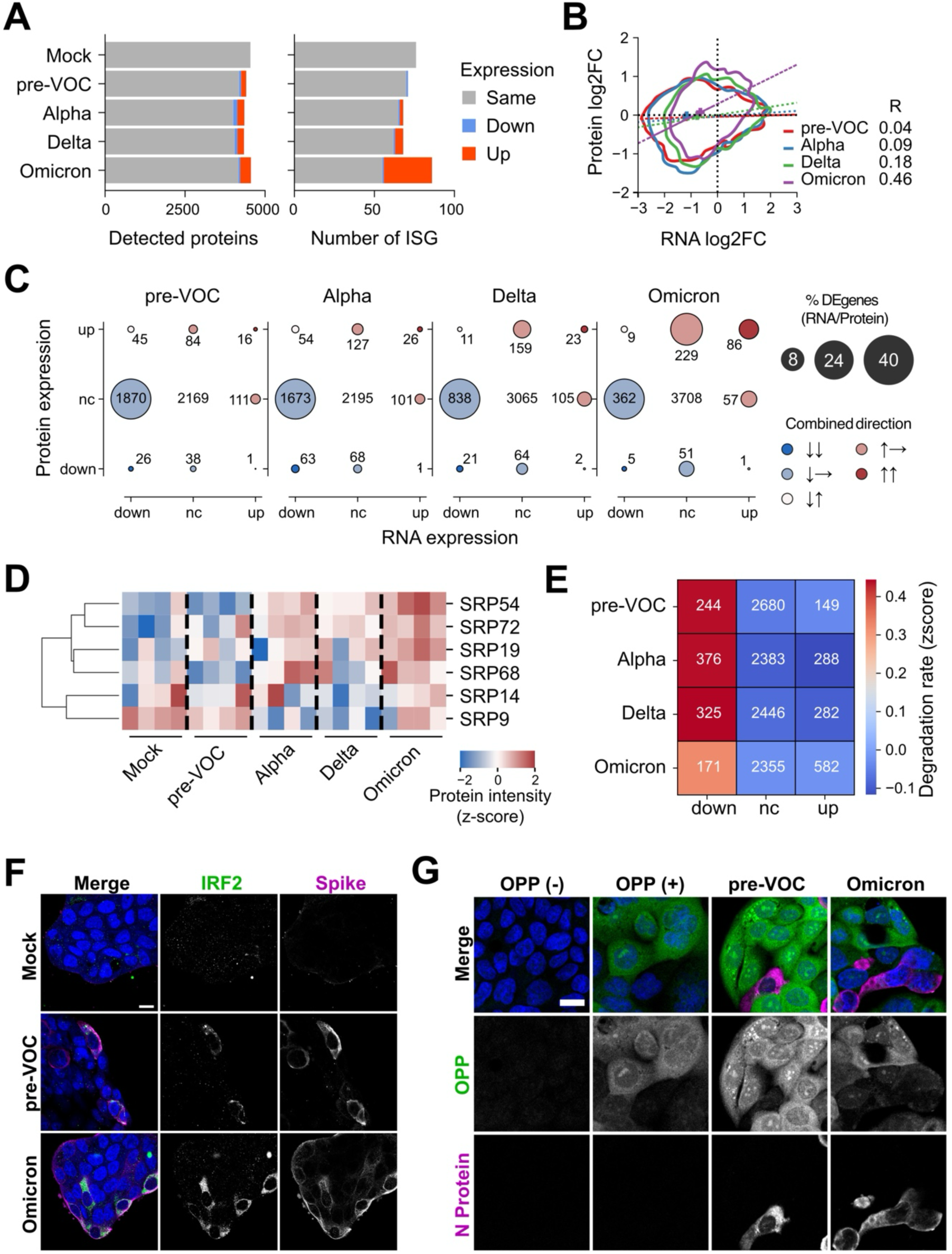
Impact of SARS-CoV-2 infection on protein expression and translation. **A)** Number of total proteins (A) or ISG (B) detected across Mock and infected conditions. Proteins significantly up- and downregulated following infection (padj < 0.05, log2-fold change > 1 or < −1) are highlighted in red and blue, respectively. **B)** Two-dimensional kernel density estimates of RNA log2 fold changes versus protein log2 fold changes for each condition (solid contours; p < 0.05). The density maximum for each condition is indicated by a cross. Dashed lines represent linear regressions, and the corresponding Pearson correlation coefficients (R) are annotated. **C)** Number of genes binned by their RNA and protein relative expression for each variant. ‘up’ and ‘down’: log2 fold-change > 1 and < −1, respectively, and adjusted p-value < 0.05. nc: not significantly changed. Numbers indicate the gene count per bin. Bubble sizes represent the percentage of differentially expressed genes, either at the RNA of protein level. Colors encode the combined direction of RNA and protein expression change (concordant up, concordant down or discordant changes). **D)** Heatmap showing the standardized degradation rate of proteins upregulated (up, padj<0.05 and fold-change>1.2), not changed (nc) or downregulated (down, padj<0.05 and fold-change<0.8) across infections, and the corresponding number of proteins. **E)** Clustermap of standardized, batch-corrected protein intensities across individual replicates (N=4) and conditions for each SRP component. **F)** Immunofluorescence staining of IRF2 and N-protein in uninfected Calu-3 cells (mock) or in cells infected with pre-VOC or Omicron viruses. **G)** Fluorescent OPP staining of infected cells combined with nucleocapsid immunofluorescence. OPP(-): negative control without OPP. OPP(+): mock cells with OPP. Scale bar represents 20 µm.

Next, to better understand protein expression alterations in infected cells, we compared protein log2 fold changes to previous large-scale assessments of protein degradation rates ^40^. This analysis revealed that downregulated proteins tend to be less stable compared to those for which expression is unchanged or increased (**Figure 6E**). Furthermore, we observed a milder correlation in cells infected by Omicron compared to pre-VOC, Alpha and Delta specimens (**Figure S5D**). These observations indicate that, in contrast to the broad perturbations in host cell RNA expression at the 48 hpi timepoint, protein expression is more resilient to infection, and the less stable proteins are the ones that exhibit the most pronounced down-regulation. GO term enrichment analysis revealed that immune response-related functional categories (e.g. defense response to virus, immune system process) were significantly enriched among genes that are upregulated both at the RNA and protein levels following Omicron infection, and to a lesser extent following Delta infection (**Figure S5E**). These terms were also enriched among genes upregulated at the RNA level, but not at the protein level, following pre-VOC and Alpha infections (**Figure S5F**). We also found that terms related to RNA metabolism, including translation components, were enriched among genes downregulated at the RNA level only or not changed (**Figure S5G-H**). Consistent with the expected induction of innate immune response proteins from our transcriptomic and proteomic profiling, we observed a striking increase in labeling for the IRF2 by IF in pre-VOC and Omicron infected cells (**Figure 6F**). Finally, to more directly visualize the impact of SARS-CoV-2 infection on mRNA translation, we next treated live pre-VOC and Omicron infected cells at 48 hpi with O-propargyl-puromycin (OPP), which allows the labeling, fluorescent tagging and visualization of nascently translated peptides ^41^. As shown in **Figure 6G**, while this labeling typically results in the generation of a robust cytoplasmic signal in mock and bystander cells, we observed a striking decrease in labeling intensity in infected cells, consistent with the expected global decrease in mRNA translation induced by SARS-CoV-2 infection ^4,42,43^.

## Discussion

In the present study, we sought to better understand the impact of SARS-CoV-2 variant infections on the RNA regulatory landscape of the human lung adenocarcinoma cell line Calu-3, via a comparative transcriptomic and proteomic profiling. Moreover, through systematic IF-based screens with a collection of RBP antibodies, we assessed the effects of infections with SARS-CoV-2 variants and abiotic immune stimulation with poly(I:C) on the subcellular distribution properties of ∼185 RBPs. As summarized in the model presented in **Figure 7**, all these conditions triggered widespread alterations of host cytoplasmic mRNAs and induced nuclear accumulation of polyA+ transcripts, polyA-binding proteins, and 7SL RNA. By contrast, the subcellular localization of most poly(I:C)-sensitive RBPs remained unchanged during SARS-CoV-2 infections. Furthermore, by contrast to early SARS-CoV-2 variants, we found that Omicron infected cells exhibited reduced viral RNA load and increased protein production of interferon stimulated genes, as well as factors associated with mRNA translation, suggesting that SARS-CoV-2 has evolved toward sustaining translation of both viral transcripts and immune response transcripts, while broadly preserving RBP subcellular distribution We observed striking differences in the amount of viral RNA produced by different SARS-CoV-2 variants, despite their similar levels of viral proteins and infection rates (**Figure 1**). Notably, Omicron BA.1 expressed substantially less viral RNA compared to previous variants. Omicron also induced a stronger immune response at the protein level (**Figure 6**), which may seem counter intuitive considering that dsRNA induced by SARS-CoV-2 infection is known to trigger the host immune response ^16^. However, at the RNA level, ISG induction was comparable across variants. It is thus possible that Omicron allows a better translation of innate immune response factors. In support to this hypothesis, we found SRP components and translation factors to be more abundant in Omicron infected cells compared to previous variants (**Figure 6**).

**Figure 7.**
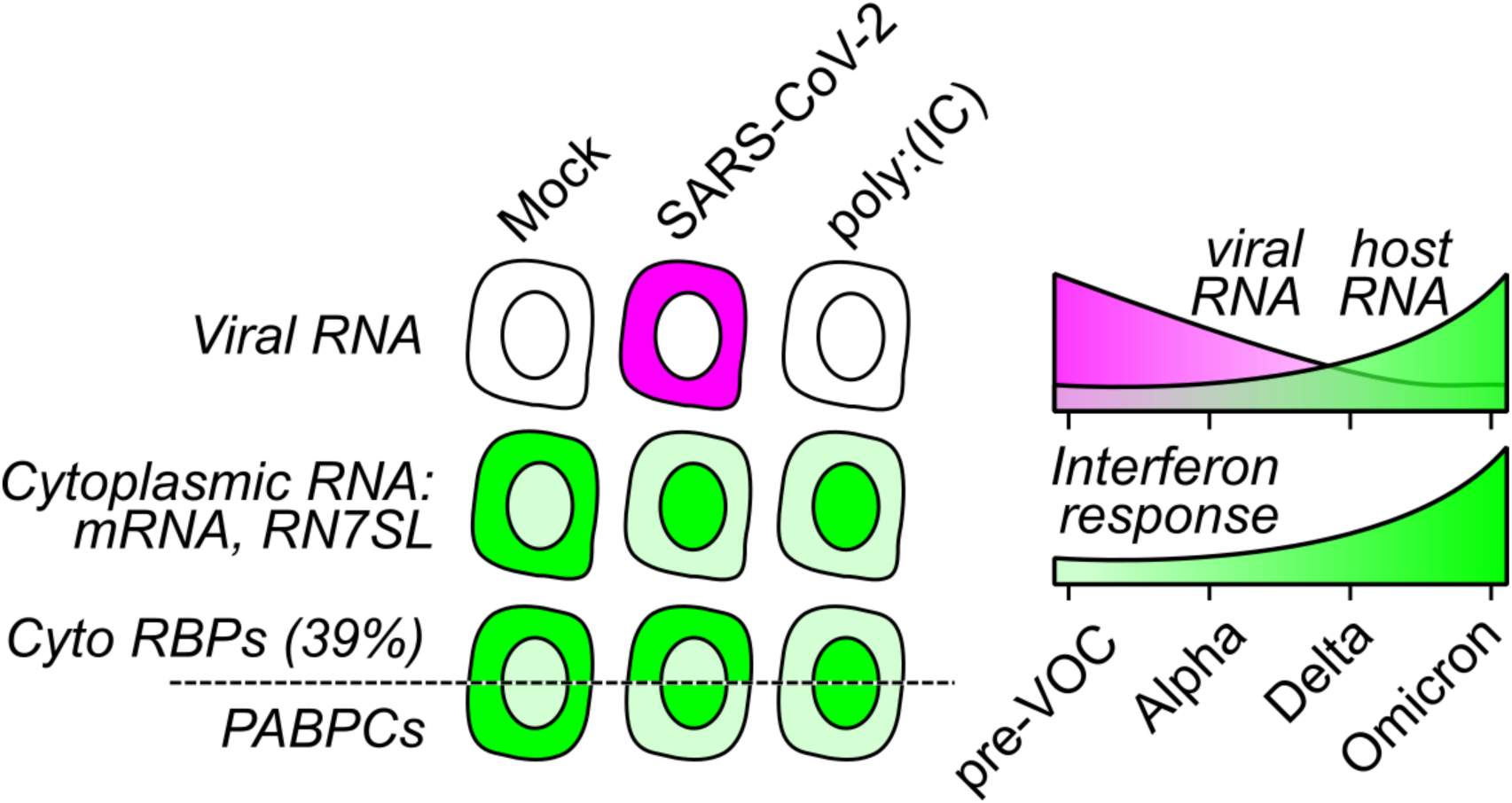
Graphical overview of the study.

Another intriguing observation from this study is the accumulation of dsRNA beyond sites of replication (**Figure 1**), i.e. sites where (-) RNA is observed. Antibodies for dsRNA (J2 or 9D5 clones) are widely used as infection markers, and it is often assumed that dsRNA forms from (-) RNA annealing to (+) RNA. In contrast, our results show that domains of (+) RNA are recognized by 9D5 antibody, in line with the previous secondary structure analysis that revealed large secondary structure elements^44^.

Our results also shed light on the pervasive remodeling of the host transcriptome and the degradation of cytoplasmic mRNAs. Unexpectedly, this major remodeling induced minimal nuclear accumulation of RBPs, except for the poly-A binding proteins. This observation contrasts with our observations from poly:(IC) stimulation, which triggered a strong nuclear translocation of several cytoplasmic RBPs. In the context of poly(I:C), previous observations based on a smaller set of RBP ^9^, led the authors to interpret it as a consequence of RBP titration by nuclear RNA. This raises the possibility that RBP subcellular localization may be driven, at least in part, by overall RNA abundance of RNA, a property that is maintained in the context of infections where the depleted host cytoplasmic mRNA pools are replace by viral counterparts. In agreement with this hypothesis, we found that poly(I:C)-sensitive RBPs were enriched with SARS-CoV-2 RNA interactors (**Figure S4**), which are enriched for RBPs predominantly involved in mRNA stability control (**Figure 4**, ^17^). It also noteworthy that among these RBPs, cytoplasmic RBPs such as DDX3X and UBAP2L have no known defined RNA binding motif ^45^, while other factors such as YBX3 are known to bind a broad range of RNA. Last, the observation that poly(A)-binding proteins are the only proteins among the ones we profiled that undergo nuclear translocation during infection also supports the contribution of RNA in the control of RBP localization. Indeed, our results indicate a pervasive reduction of the abundance of cytoplasmic, poly(A) mRNAs. In addition, viral RNA is abundant in infected cells, but possesses shorter poly(A) tails compared to host mRNAs ^17^. Consequently, the nuclear translocation of poly(A)-binding proteins may result from reduced availability of cytoplasmic poly(A) binding sites. RBP localization may thus be opportunistically driven by RNA binding site availability. Beyond its role as a genetic template, viral RNA could therefore act as a scaffold to maintain cytoplasmic RBP concentration in a range that permits replication. In addition, SARS-CoV-2 has been shown to interfere with nucleocytoplasmic trafficking ^4,46^ and may block nuclear import of RBPs upon cytoplasmic RNA degradation ^16,47^.

## Material & Methods

### Cell culture

Human lung epithelial calu-3 cells and A549 cells were maintained in EMEM supplemented with 10% FBS, and DMEM supplemented with 5% FBS and 1% pen/strep, respectively, and grown in a humidified incubator at 37°C with 5% CO2.

### Preparation of SARS-CoV-2 stocks and cell infections

The ancestral SARS-CoV-2 variant used in this study is LSPQ1 (B1 lineage, GISAID: EPI_ISL_535728), isolated in Quebec by the Laboratoire de Santé Publique du Québec (LSPQ). This isolate is similar to the Wuhan strain (PMID: 35812400). The Alpha variant (B.1.1.7, GISAID: EPI_ISL_683466) was obtained from BEI Resources. The Delta variant (B.1.617.2, ID: L00414346/COV2021-0248 3384976) was obtained from the National Microbiology Laboratory (Winnipeg, Canada). LSPQ1, Alpha (B.1.1.7), and Delta (B.1.617.2) variants were amplified and titrated in Vero E6 cells using a previously described plaque assay (PMID: 39804090). The Omicron BA.1 (B.1.529, ID: L00423388) variant, obtained from LSPQ, was amplified once in Calu-3 cells and titrated in Vero-TMPRSS2 by plaque assay. Other Omicron isolates (BA.1, BA.5., JN.1) were obtained from BEI Resources and similarly amplified and titrated. Calu-3 cells were infected with LSPQ1, Alpha, Delta, and Omicron for 48 h at MOIs ranging from 0.001 to 0.5 to obtain similar levels of nucleocapsid-expressing cells, as determined by flow cytometry as in (PMID: 39804090).

### Poly(I:C) transfection

Cells were seeded at 4,000 cells per well in 96-well plates and allowed to adhere overnight. Poly(I:C) (InvivoGen; stock 0.01 µg µL⁻¹ in nuclease-free water) was delivered to cells using jetPRIME (Polyplus) according to the manufacturer’s recommendations with minor modifications. For each well, 0.04 µg poly(I:C) was diluted in 40 µL jetPRIME buffer and vortexed briefly. jetPRIME reagent was added at a ratio of 2 µL reagent per µg poly(I:C) (0.08 µL per well), followed by a 1-s vortex. The mixture was incubated for 10 min at room temperature, then adjusted to a final volume of 200 µL using pre-warmed DMEM containing 5% FBS and 1% penicillin–streptomycin. Before transfection, culture medium was aspirated, and cells were washed once with PBS. The poly(I:C)–jetPRIME mixture was added directly to the wells and cells were incubated for 1 h at 37 °C. Control wells received an equivalent volume of nuclease-free water in place of poly(I:C). Following transfection, cells were washed three times with PBS and returned to complete medium for a 5-h recovery period at 37 °C.

### Immunofluoresence and FISH

The protocol was adapted from a previous study ^15^. Cells were rinsed in PBS and fixed in 3.7% formaldehyde for 4 hours, washed in PBS and permeabilized in PBS and 0.1% Triton X-100 (PBS-T). Primary antibodies were diluted in PBS-T and incubated with samples for 1 hours at room temperature. Dilutions were 1:200 for 1 mg/mL and 1:100 for 0.2 mg/mL stock concentration. After three washes in PBS-T, samples were incubated for 1 hour with Alexa Fluor 488 or Alexa Fluor 647-conjugated secondary antibody (Thermo Fisher Scientific) diluted in PBS-T at 1:1000. After three PBS-T washes and two PBS washes, samples were either stained with DAPI or post-fixed in 3.7% formaldehyde for 10 min for subsequent FISH staining. For IF-FISH, samples were washed twice in PBS after post-fixation and equilibrated in 10% formamide, 2X saline-sodium citrate (SSC 2X). SARS-CoV-2 specific probes (final concentration: 4 µM) were annealed to Cy-3 or Cy-5 labeled complementary flap oligonucleotides (final concentration: 5 µM) in 100 mM NaCl, 50 mM Tris-HCl pH7.5, 10 mM MgCl2, 1 mM DTT by heating at 65°C for 5 minutes then incubating at 35°C for 15 minutes. Samples were incubated overnight at 37°C with the fluorescent probe duplexes in 10% formamide, 2X SSC, 1% dextran sulfate and 0.4 U/µL RNAseOUT (Invitrogen). After hybridization, samples were washed twice in 10% formamide, 2X SSC for 30 min at 37°C followed by two PBS washes. Nuclei were stained with 1 µg/mL DAPI for 5 min and rinsed with PBS. Images were acquired using either an automated Observer 7 microscope (Zeiss) equipped with a 40X dry objective or a CSU-1 spinning disk (Yokogawa) confocal microscope (Zeiss) equipped with a 63X oil-immersion objective.

O-propargyl-puromycin (OPP) incorporation was assessed using the Click-iT Plus OPP Protein Synthesis Assay Kit (Molecular Probes, C10456). Cells were incubated with 20 µM OPP for 30 min prior to fixation. Samples were immunolabeled with an anti-nucleocapsid primary antibody (Sinobiological, 40143-MM05-100) and an Alexa Fluor 647-conjugated secondary antibody as previously described After. PBS wash, the Click-iT reaction was performed according to the manufacturer’s instructions.

### RNA preparation for RT-qPCR and sequencing

Calu-3 were seeded in 12-well plates and infected using 3 wells per variant and each well was processed independently. After 48h, RNA was extracted using TRIzol reagent (Thermo Fisher, 15596018) following the manufacturer’s recommendations. RNA purification was purified using the RNA Clean & Concentrator-5 system (Zymo Research, R1013) and in-column DNaseI treatment. The RNA was eluted in nuclease-free water and quantified by Nanodrop.

### RT-qPCR

Complementary DNA was generated using random hexamers, oligo dT and superscript III reverse transcriptase (Invitrogen) using 500ng of total RNA from each sample. Quantitative PCR (qPCR) analysis was performed with gene-specific primer pairs, using the Power up™ SYBR® green Master Mix (Applied Biosystems) on an ABI ViiA7 instrument (Life Technologies, Inc). Each reaction was carried out in duplicates and three independent experiments were run. 18S rRNA was used as an internal control.

### RNA sequencing

Calu-3 were seeded in 12-well plates and infected using 3 wells per variant and each well was processed independently. After 48h, RNA was extracted using TRIzol reagent (Thermo Fisher, 15596018) following the manufacturer’s recommendations. RNA purification was purified using the RNA Clean & Concentrator-5 system (Zymo Research, R1013) and in-column DNaseI treatment. The RNA was eluted in nuclease-free water and quantified by Nanodrop and Bioanalyzer. After the extraction of RNAs, the samples were ribo-depleted followed by library preparation with the KAPA RNA HyperPrep Kit with RiboErase (HMR) (Roche, 8098140702). Libraries were PCR-amplified, quantified by qPCR and loaded at equimolar concentrations for sequencing on the Novaseq platform with the kit Novaseq S4 (Illumina) at a coverage of 100M paired-end reads per sample.

### RNAseq bioinformatic analyses

The FASTQ files were trimmed to remove sequencing adapters and positions with poor quality using Trimmomatic (PMID: 24695404) v0.39 with following parameters:-phred33 ILLUMINACLIP:TruSeq3-PE.fa:2:12:10:8:true TRAILING:30 LEADING:30 MINLEN:3. The quality of the reads was assessed before and after trimming with FastQC (https://www.bioinformatics.babraham.ac.uk/projects/fastqc/) v0.11.9. The viral genome built based on Ensembl (PMID: 36318249) Sars_cov_2.ASM985889v3 annotation was combined to the human genome (GRCh38.p109 from Ensembl) and was used to build the indexes for the alignment step using STAR (PMID: 23104886) v2.7.11b with the option --runMode genomeGenerate --sjdbOverhang 149. The trimmed FASTQ files were aligned to their respective index with STAR --runMode alignReads --outFilterType Normal --outFilterMultimapNmax 100 --winAnchorMultimapNmax 100. The human gene quantification was done using CoCo (PMID: 31141144) v1.0.0 using coco ca with default parameters and coco cc with -c both -s 2 -p. The differential expression analysis was assessed by DESeq2 (PMID: 25516281) v1.42.0. For the visualisation of the alignments on the genomic tracks, the BAM files were filtered to only keep primary alignments to the human genome with SAMtools (PMID: 19505943) v1.20. The BAM files were sorted and indexed with samtools sort and samtools index and filtered using samtools view -F 256, removing alignments to the viral genomes using grep -vw MN908947.3. The CPM normalised bedgraphs were generated with deepTools (PMID: 27079975) v3.5.5 bamCoverage command with the options --outFileFormat bedgraph -bs 1 --normalizeUsing CPM. The resulting bedgraphs were averaged to get one file per condition using a custom script.

### Protein preparation for mass spectrometry

Calu-3 were seeded in 12-well plates and infected using 4 wells per variant. Each well was processed independently. After 48h, cells were rinced with PBS and lysed in 6M guanidium hydrochloride and 100 mM Tris-HCl pH8. Samples were heat-inactivated at 95°C for 5 minutes. Protein samples were then analyzed by liquid chromatography-tandem mass spectrometry (LC-MS/MS) on the Easy-nLC II nano-flow liquid chromatography system (Proxeon Biosystems), coupled with an Orbitrap Fusion mass spectrometer (Thermo Scientific) via a Nanospray Flex Ion Source (Thermo Scientific).

### Proteomics bioinformatic analyses

Raw mass spectrometry files were transformed to the .mzML format using ProteoWizard’s MSConvert tool version 3.0.22055 ^48^ and analyzed by FragPipe (version 21.1) ^49^. To prevent MBR (Match Between Runs) biases, searches were performed separately for each of the Mock or the mutants using the LFQ-MBR default workflow settings. Searches were executed against the Uniprot human non-redundant reference proteome (UP000005640 2022_12 release) ^49^, which was supplemented with the non-redundant SARS-Cov-2 proteome (UP000464024 2024_02 release), the universal contaminant library ^50^ and reverse decoy sequences. The MSFragger parameters were set to default with trypsin specificity (two missed cleavage sites allowed), oxidation (M) and N-terminal acetylation as variable modifications, and cysteine carbamidomethyl as a fix modification. Output files generated by FragPipe (“msstats.csv”), one for each of the five conditions, were read and assembled in R (r-project.org). Once assembled, reproducibility and batch effects among biological replicates were evaluated in R by generating multidimensional scaling (MDS) plots with the limma package (Ritchie et al, 2015). The data was adjusted for a batch effect associated with the day of mass spectrometry (MS) acquisition. This was done by using the removeBatchEffect function from the limma R package. Once the data was corrected, it was converted into the MSstats format^51^ with the FragPipetoMSstatsFormat function with the following parameters: useUniquePeptide = TRUE, removeFewMeasurements = TRUE, removeProtein_with1Feature = FALSE, summaryforMultipleRows = max. The MS data were cleaned, imputed, ‘equalizeMedians’ normalized, and summarized using the robust Tukey’s median polish estimation method by applying the ‘dataProcess’ function from the MSstats package. Statistical contrasts were next performed using MSstats’ groupComparison function. For each statistical contrast, if the intensity of either the control or the condition was missing, the log2 fold change was imputed with the condition’s log2 intensity or the negative log2 control intensity. Proteins displaying an adjusted p-value ≤ 0.05 and a Log2 fold change ≤ −1 or ≥ 1 were considered statistically significant.

## Acknowledgments

The authors are grateful for the support from colleagues at the IRCM microscopy (Dominic Fillon and team), molecular biology (Sarah Boissel and team), and mass spectrometry (Denis Faubert and Josée Champagne) platforms. G.D.B. was funded by a postdoctoral fellowship from the Fonds de Recherche du Québec Santé (FRQS), as well as a scholarship from the IRCM foundation. E.L. is an FRQS distinguished research scholar. This work was supported by grants to E.L. from CIHR, the Québec Ministry of Economy, Innovation and Energy (MEIE), as well as an FRQS joint research chair in AI & Health research. É.A.C. is the recipient of the IRCM-Université de Montréal Chair of Excellence in HIV Research.

---

**Figure S1.**
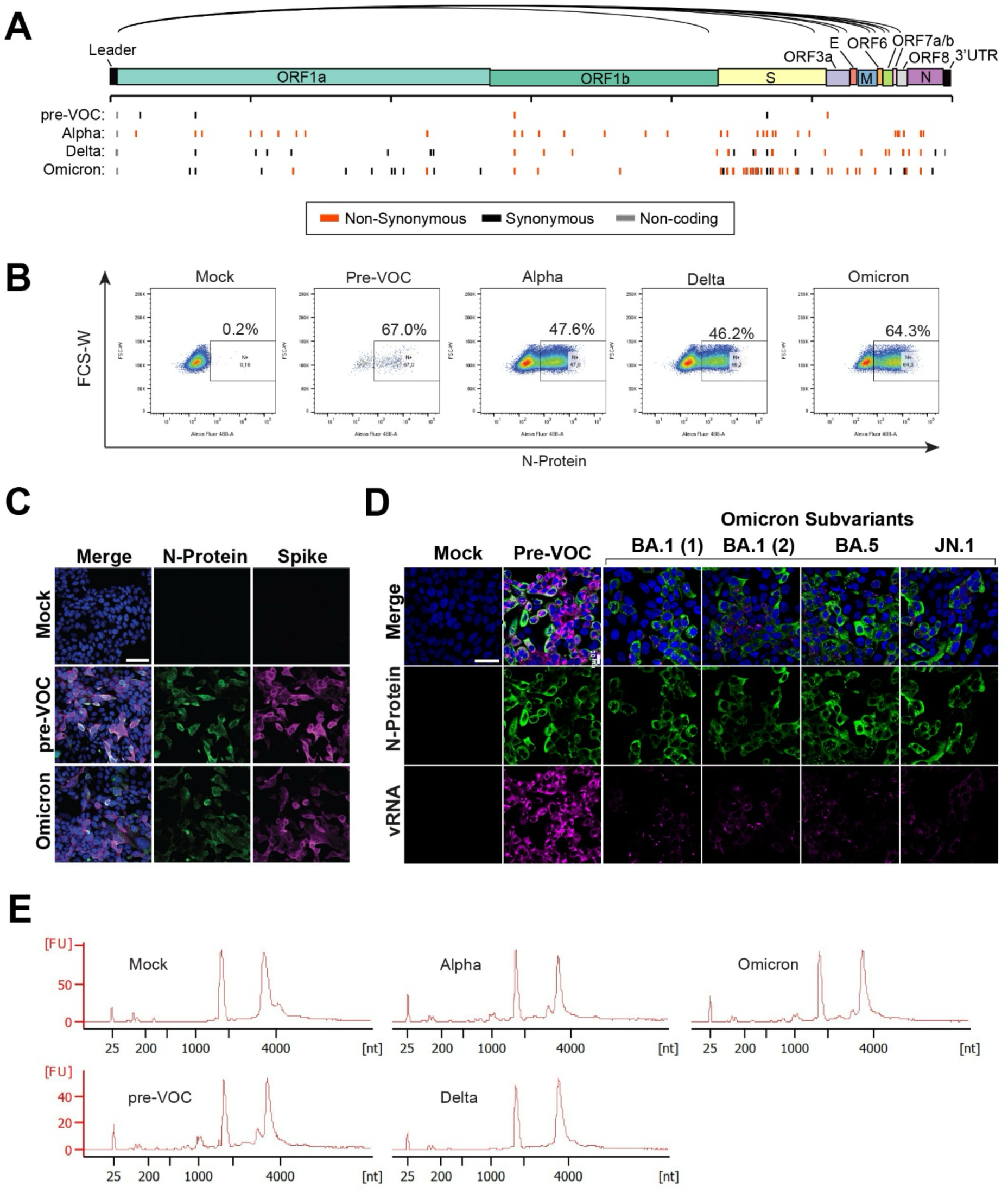
Comparative features of SARS-CoV-2 variants interrogated in this study. **A)** Architecture of the SARS-CoV-2 genome, with black lines indicating canonical junctions yielding subgenomic RNAs. Mutations in pre-VOC, Alpha, Delta and Omicron BA.1 isolates are shown relative to the SARS-CoV-2 reference sequence (NC_045512.2). Non-coding, synonymous and non-synonymous mutations are highlighted in grey, black and red, respectively. **B)** Representative flow-cytometry plots showing nucleocapsid intensity at 48 hpi. **C)** Immunofluorescence staining of infected Calu-3 cells with an antibody against SARS-COV-2 nucleocapsid and spike proteins. Scale bars: 40 µm. **D)** Immunofluorescence staining for nucleocapsid protein combined with viral RNA FISH in pre-VOC, Omicron BA.1 and additional Omicron isolates corresponding to BA.1, BA.5 and JN.1 subvariants. **E)** Representative bioanalyzer traces showing similar levels of 18S and 28S rRNA between infected and mock samples.

**Figure S2.**
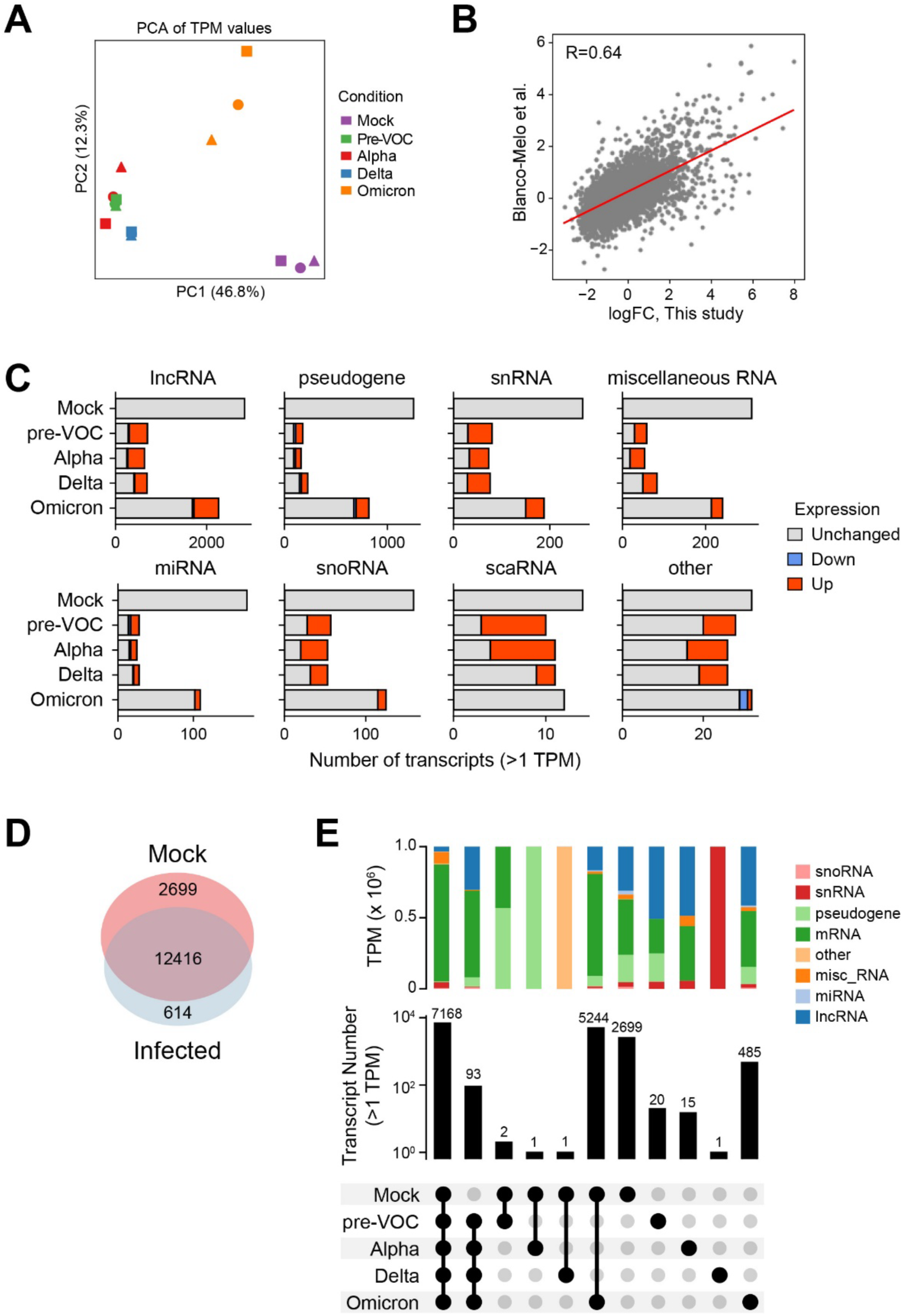
Transcriptomic features of Calu-3 cells infected with different SARS-CoV-2 variants. **A)** Principal component analysis (PCA) of log2-transformed host TPM values across infection conditions and replicates. **B)** Comparison of gene expression log2 fold change between Blanco-Melo et al. (2020) and this study (pre-VOC condition, TPM>1). The red line represents a linear regression fit, with the Pearson correlation coefficient (R) indicated. **C)** Number of detected transcripts (TPM>1) across various RNA biotypes, with the number of over- or under-represented transcripts indicated in red and blue, respectively, while gray indicates transcripts for which the representation is unchanged relative to Mock.**<u>D)</u>** Venn diagram showing transcripts detected exclusively in mock samples versus those detected in any infected sample (TPM>1). **E)** UpSet plot showing the intersection sizes of transcript detected (TPM>1) across samples (bottom) and the proportion of sequencing reads attributable to different biotypes within each intersection (top).

**Figure S3.**
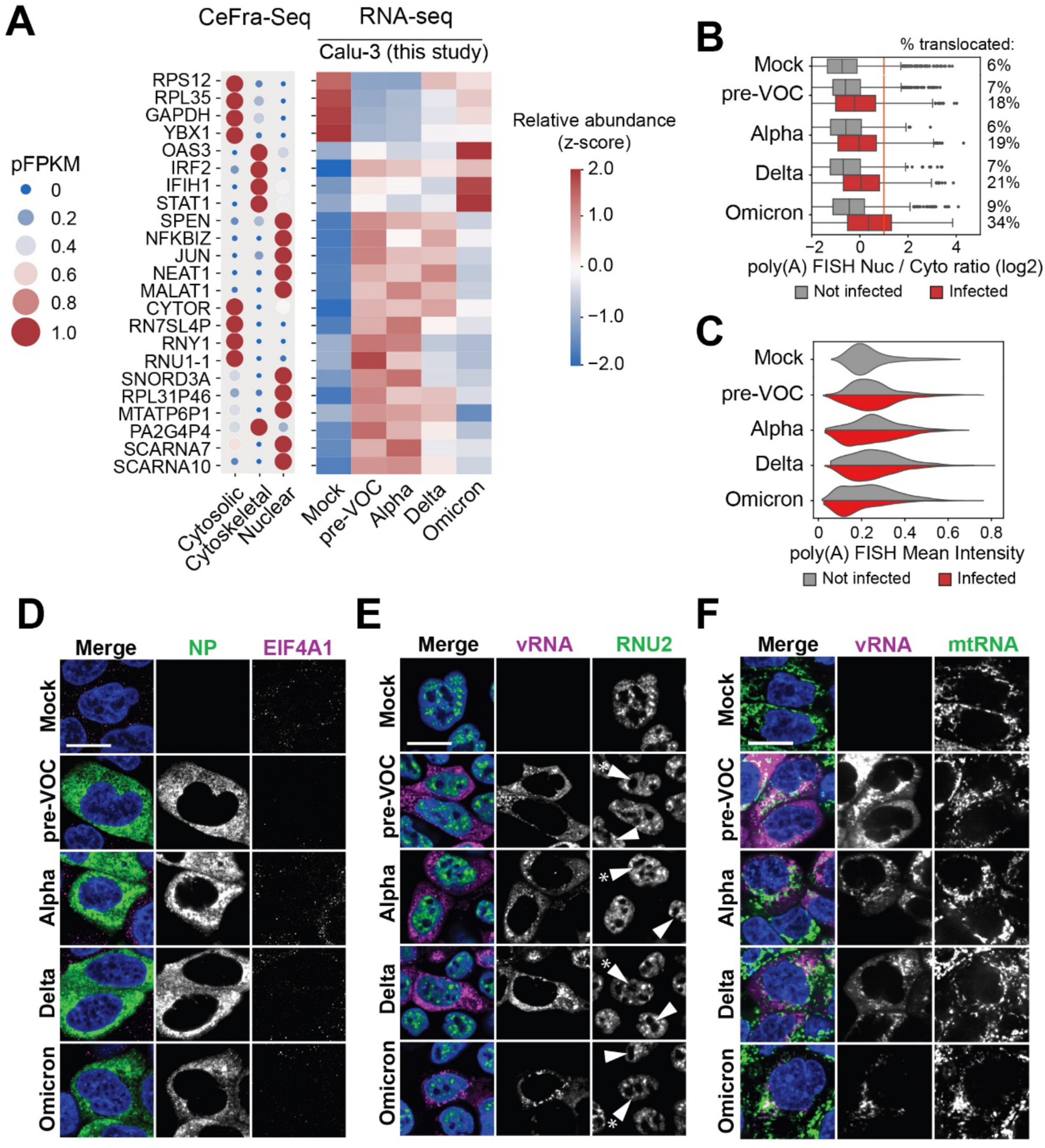
RNA subcellular localization features of SARS-CoV-2 infected cells. **A)** Examples of specific RNA transcripts exhibiting asymmetric localization patterns within CeFra-seq datasets (left panel), compared to their respective expression (in TPMs) across in mock or infected Calu-3 cells. **B)** Boxplot showing the log2 ratio of nuclear over cytoplasmic mean intensity in uninfected (gray) vs infected (red). **C)** Violinplot showing polyA (+) RNA mean FISH intensity in uninfected (gray) vs infected (red) cells. The percentage of cells with nuclear enriched polyA (+) RNA (log2 ratio >1) is indicated on the right. **D-F)** FISH analyses of EIF4A1 mRNA (**D**), the RNU2 snRNA (**E**), mitochondrial heavy and light chain RNA/mtRNA (**F**) in conjunction with N-protein immunostaining (**D**) or viral RNA FISH (**E,F**). Scale bars: 20 µm.

**Figure S4.**
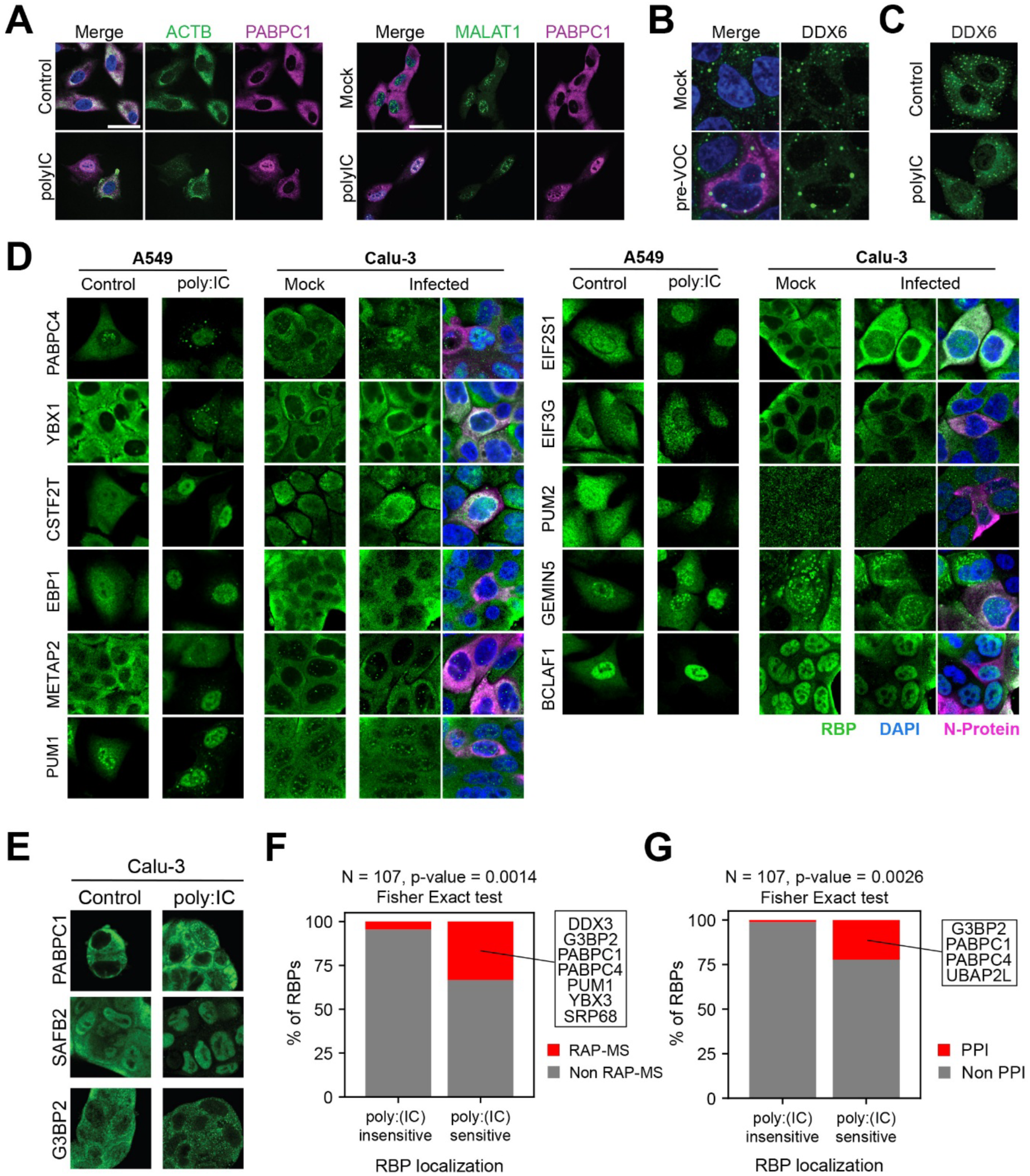
RPB subcellular localization features in poly: IC treated versus infected cells. **A-B)** PABPC1 immunofluorescence along with RNA FISH patterns of *ACTB* mRNA (left) and *MALAT-1* lncRNA (right) in untreated or poly:IC treated A549 cells. **B-C)** DDX6 immunofluorescence in Mock vs pre-VOC infected Calu-3 (B) or untreated vs poly:IC treated A549 cells. **D)** Immunostaining patterns of RBPs with perturbed subcellular localization upon poly:IC transfection in untreated or poly:IC treated A549, or in mock or pre-VOC infected Calu-3 cells. **C)** Immunofluorescence of PABPC1, SAFB2 and G3BP1 in untreated or poly:IC transfected Calu-3 cells. **F)** Percentage of poly:IC-sensitive and poly:IC-insensitive RBPs annotated as SARS-CoV-2 RNA interactors (left, ^36^) or SARS-CoV-2 protein interactors (right, ^37^). **G)**

**Figure S5.**
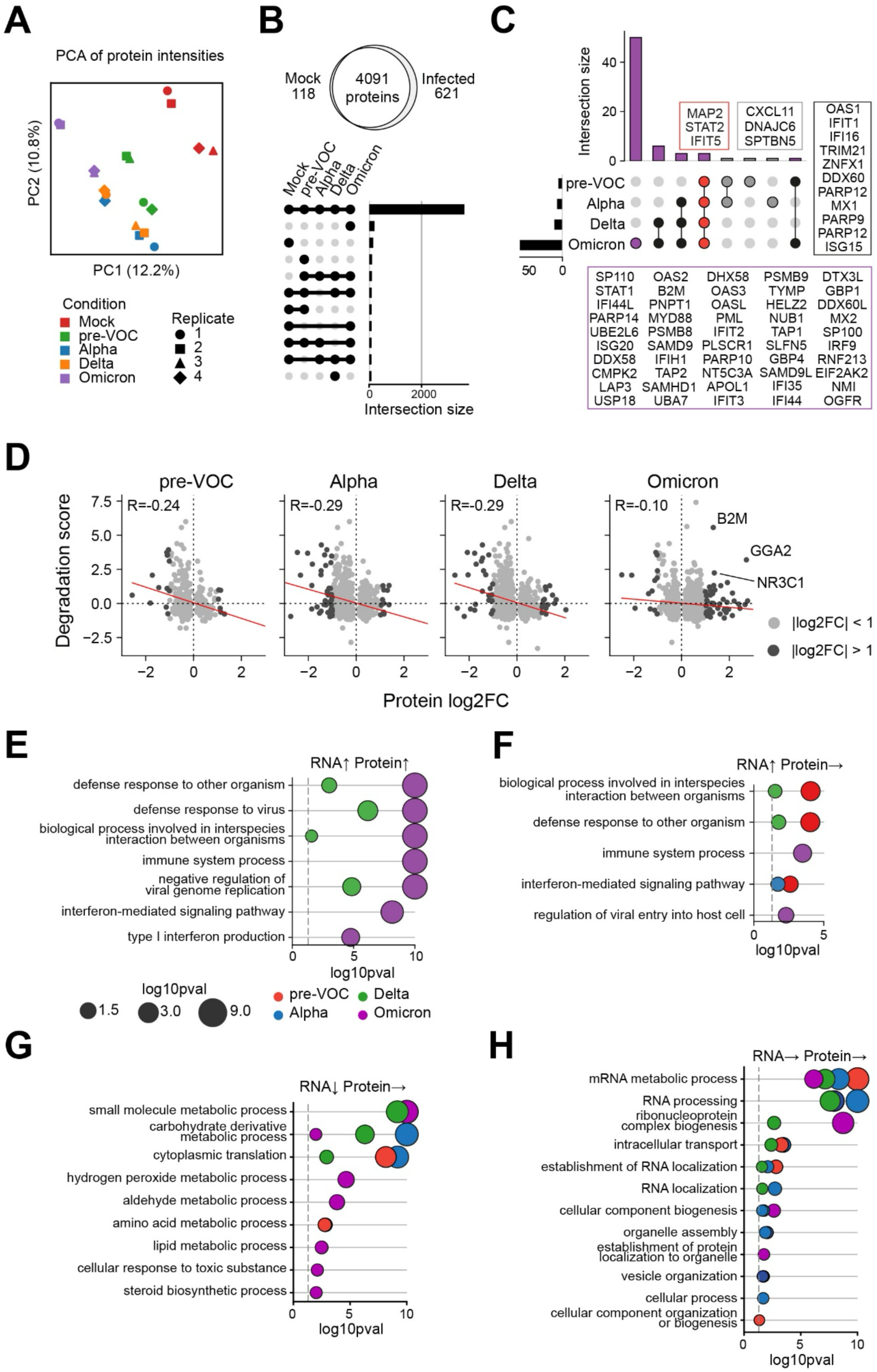
Translation of immune response genes is more efficient in Omicron. **A)** Principal component analysis (PCA) of log2-transformed host protein intensities across infection conditions and replicates. **B)** Venn diagram and upset plot of the total number of proteins detected in mock versus infected Calu-3 cells. **C)** Upsetplots showing upregulated ISG proteins (log2 fold-change > 1, p-value < 0.05) across infections. Categories including Omicron are indicated in purple while categories excluding Omicron are in grey, with the corresponding genes indicated in colored boxes. Genes highlighted in red are upregulated across all variants. Genes in bold are upregulated in at least two variants, while other are upregulated only following Omicron infection. **D)** Scatterplot showing protein degradation rates (z-scores; data form PMID: 24045637) plotted against protein log2 fold changes (this study). Only proteins with significant expression changes (adjusted p-value < 0.05 and log2 batch-corrected intensity > 10) are included. Red lines indicate linear regression fits, and the corresponding Pearson correlation coefficient (R) are annotated. Proteins with log2 fold-change > 1 or < −1 are highlighted in black while other proteins are in grey. Selected proteins with high degradation rates (z-score > 2) and strong up-regulation (log2FC > 1) are labeled. **E-H)** Bubble plots showing significantly enriched GO biological process terms for genes upregulated at both the RNA and protein level (**E**), upregulated at the RNA level and not changed at the protein level (**F**), downregulated at the RNA level and not changed at the protein level (**G**) or not changed at both levels (**H**). Variants are indicated by distinct bubble colors. Bubble sizes represent the enrichment significance (log10 p-value), with values capped at 10. Dashed lines mark the significant threshold (p-value = 0.05).

**Figure S6.**
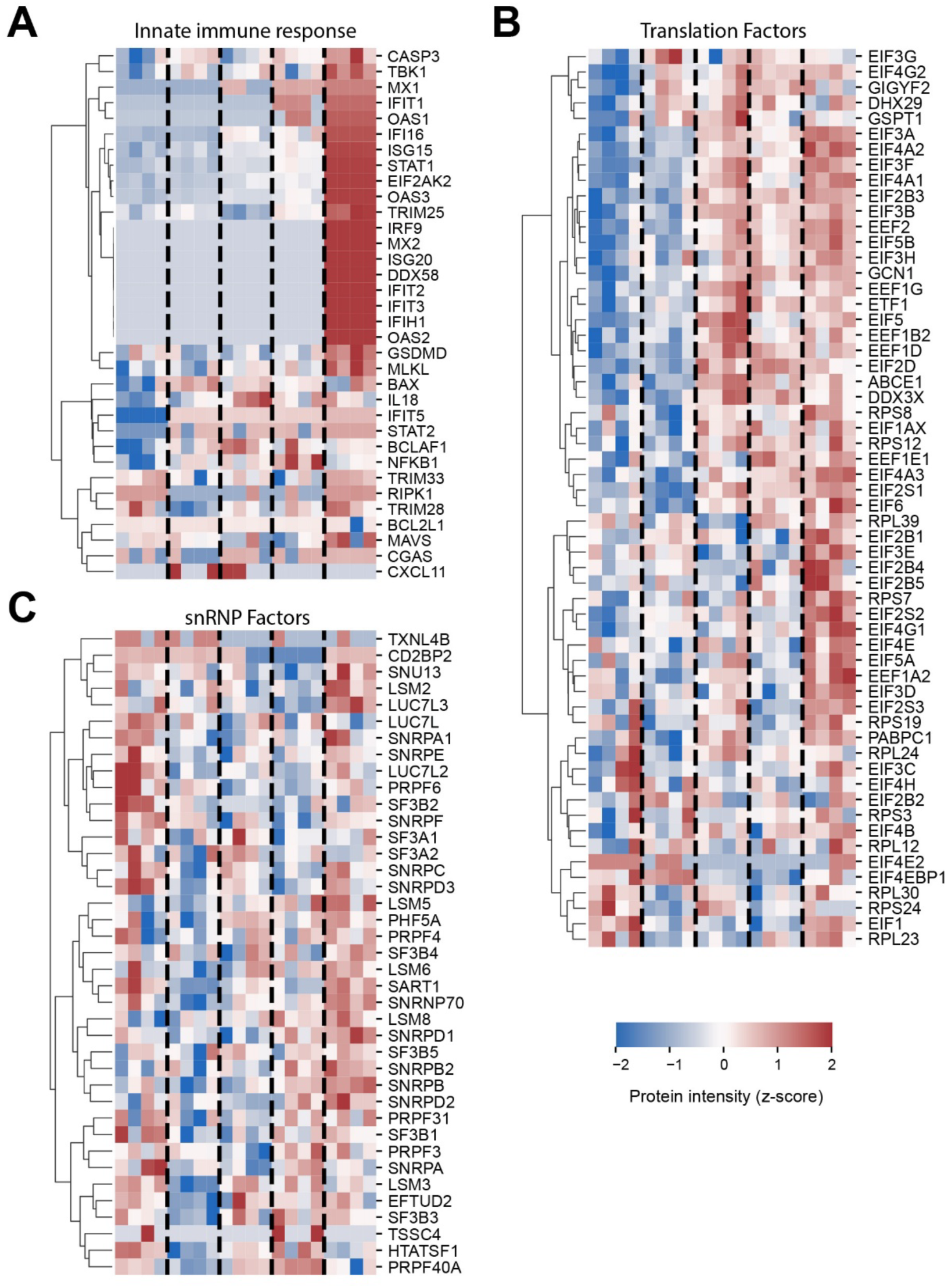
Comparative intensity signatures of proteins associated with the innate immune response, splicing and translation. **A-C)** Clustermaps of standardized, batch-corrected intensities across individual replicates (N=4) and conditions for selected immune response factors (**A**; including PRRs, adaptors, ISGs, cytokines and cell-death pathway components), translation factors (**B**; including ribosomal proteins, initiation, elongation, termination, and accessory factors) and snRNP components (**C**; including core components and U2 snRNP-specific components).

